# FIST-*n*D: A tool for *n*-dimensional spatial transcriptomics data imputation via graph-regularized tensor completion

**DOI:** 10.1101/2022.10.12.511928

**Authors:** Thomas Karl Atkins, Tianci Song, Rui Kuang

## Abstract

Functional interpretation of spatial transcriptomics data usually requires non-trivial pre-processing steps and other supporting data in the analysis due to the high sparsity and incompleteness of spatial RNA profiling, especially in 3D constructions. As a solution, we present a new software tool FIST-*n*D, Fast Imputation of Spatially-resolved transcriptomes by graph-regularized Tensor completion in *n*-Dimensions for imputing 3D as well as 2D spatial transcriptomics data. FIST-*n*D is implemented based on a novel graph-regularized tensor decomposition method, which imputes spatial gene expression data using 4-way high-order tensor structure and relations in spatial and gene functional graphs. The implementation, accelerated by GPU or multicore parallel computing, can efficiently impute high-resolution 3D spatial transcriptomics data within a few minutes. The experiments on three 3D Spatial Transcriptomics datasets and one 3D high-resolution Stereo-seq dataset confirm the high accuracy of the imputation by FIST-*n*D and demonstrate that the imputed spatial transcriptomes provide a more complete gene expression landscape for downstream analyses such as spatial gene expression clustering and visualizations.

## Background

To understand how the composition and arrangement of cells in tissues contribute to their functional role has long been one of the primary goals in biological research (Vesalius, 1543). One way to accomplish this is to measure the presence of RNA in a tissue. As mRNA acts as a precursor to proteins, measuring the levels of mRNA in a tissue can give insight into the biological processes occuring in that tissue. While RNA sequencing (RNA-seq) and single-cell RNA sequencing (scRNA-seq) allow researchers to quantify the transcriptome in bulk tissue samples as well as single cells, the spatial variation in expression patterns is still needed for a complete mapping of the cells’ organization in a tissue (Miller et al., 2021). More recently developed in-situ capturing (ISC) methods perform RNA sequencing of the whole transcriptome with positional barcodes in a spatial genomic array aligned to locations on the tissue, including Spatial Transcriptomics (ST) (Ståhl et al., 2016) (commercialized as 10x Genomics Visium (10x Genomics, 2019)), higher resolution Slide-seq (Rodriques et al., 2019), high-definition spatial transcriptomics (HDST) (Vickovic et al., 2019) and Spatiotemporal enhanced REsolution omics-sequencing (Stereo-seq) (Chen et al., 2022). These ISC-based methods capture two dimensional gene expression profiles from a tissue section attached to a slide, where RNA is hybridized to primers that contain both a spatial barcode and unique molecular identifiers (UMIs) that, when prepared into a DNA library, allows researchers to determine both the location and the transcript where a read originated. From this, a “map” of RNA expression across a tissue slice can be constructed to give the levels of RNA expression at every spot on the slide.

Though current spatial transcriptomics platforms collect data from a two-dimensional tissue slice, parallel adjacent tissue slices can be combined to create a three dimensional gene expression dataset (Asp et al., 2019; Ortiz et al., 2020; Vickovic et al., 2022; Wang et al., 2022). Several methods have been developed to align these parallel slices in 3D space, either using histological imaging or similarity of expression profiles (Bergenstråhle et al., 2020; Zeira et al., 2022). The 3D construction is critical for visualizing and understanding gene expression patterns in complete spatial arrangement in 3D such as those in Drosophila embryos (Wang et al., 2022)

Due to the technical limitations of ISC such as low RNA capture efficiency (Asp et al., 2020) and failure in RNA fixation and permeabilization in some tissue regions (Li et al., 2021), integrative analysis of the 2D profiling of multiple adjacent slices of a tissue to reconstruct the gene expressions in ℝ^3^ is still a new challenge. To facilitate the analysis of the 3D gene expressions, we present FIST-*n*D (Fast Imputation of Spatially-resolved transcriptomes by Graph-regularized Tensor decomposition in *n*-Dimensions), a novel method of imputing the 3D data that incorporates spatial information and gene functional relations in a protein-protein interaction (PPI) network. FIST-*n*D is an extension of an earlier ST 2D imputation method, FIST (Li et al., 2021) with an extended 3D formulation and implementation to be applied on stacked slice data. While the original FIST completes a 3-way sparse tensor in genes and the (*x, y*) spatial coordinates, FIST-*n*D is a generalization to include the 3D spatial coordinates (*x, y, z*) and the genes in a 4-way sparse tensor. FIST-*n*D discretizes the input spatial gene expression data with continuous spatial coordinates into a tensor by a voxelization process. Then, graph-regularized tensor completion is applied to the 4D sparse tensor regularized by a spatial graph and protein-protein interaction network (PPI). Finally, the output tensor decomposition can be directly analyzed for understanding the 3D gene expression patterns or alternatively, converted to original spatial expressions through a process of multilinear interpolation. Experiments on four publicly available datasets finds that FIST-*n*D is more accurate than other methods, and that the imputation with FIST-*n*D leads to increased performance on downstream analysis tasks. We also provide an Python command line implementation of the method that is easy-to-use.

Many software pipelines have been developed for analysis of spatial transcriptomic data. STUtility is a pipeline that takes 10X Visium data and standardizes alignment of sections with annotation and visualization features (Bergenstråhle et al., 2020). Squidpy integrates multiple types of spatial data to compute a spatial graph for analysis of spatial patterns (Palla et al., 2022). Giotto is a tool that clusters spatial data and performs enrichment analyses (Dries et al., 2020). However, none of these tools adopts the imputation strategy prior to their functional analysis, limiting the information gained from the spatial data. Because modern high-throughput and transcriptome-wide spatial and single-cell sequencing technologies generate zero-inflated data, many of the zero values in the final count matrices are not reflective of actual biological zeros, but rather arise as a result of sampling biases (Jiang et al., 2022). Imputation is a strategy that has the potential to improve the functional analysis of the data with more truthful RNA expressions (Li et al., 2021), which is more eminent in 3D spatial gene expressions. So far, most existing imputation algorithms have primarily focused on imputation and dimension reduction of single-cell RNA-seq (scRNA-seq) data, rather than spatial transcriptome data. One such method, DeepImpute, imputes scRNA-seq data by splitting genes into related blocks for imputation with a sub-neural network (Arisdakessian et al., 2019). Another popular imputation technique for scRNA-seq data, MAGIC constructs a Markov distance matrix between cells, which can be used to impute the original data with a smoothing step (van Dijk et al., 2018). However, these scRNA-seq methods do not utilize information about spatial relationships between cells, and as such ignore the spatial context beneficial to the imputation. Our method, FIST-*n*D outperforms these methods by incorporating the spatial information into the imputation algorithm.

## Results

### Overview of FIST-*n*D

FIST-*n*D (Fast Imputation of Spatially-resolved transcriptomes by graph-regularized Tensor completion in *n*-Dimensions) is an extension of our earlier ST imputation method, FIST, which minimizes an objective function of graph-regularized tensor completion over the gene expression tensor and a tensor product graph of the spatial chain graphs of each spatial axis and the PPI network (Li et al., 2021). The complete mathematical definitions and the multiplicative updating algorithm are reviewed in the **Methods** section. FIST-*n*D generalizes FIST for *n*-dimensional tensor and the matched higher-order graph such that stacked 2D slice spatial gene expression data can be analyzed in 3D. From here on out, we refer to FIST-*n*D as FIST as well, as FIST-*n*D applied to two-dimensional data is identical to the original method. The FIST-*n*D pipeline is detailed in Figure 1. First, FIST takes in continuous spatial gene expression data as input and performs a voxelization process to convert the data into a discrete tensor where the first dimension corresponds to genes and all subsequent dimensions correspond to the spatial dimensions (x,y,z) of the data. In the tensor completion step, the objective function is constructed to minimize the distance between the original and imputed tensors. The objective function is also regularized by a product graph of the spatial chain graphs and protein-protein interaction network (PPI). The optimization algorithm is implemented by either a Python script (optimized for GPU computing) or a MATLAB script (optimized for multicore performance). Finally, the learned canonical polyadic decomposition (CPD) form of the tensor can be used to construct the full tensor for conversion to the original data format through a process of multilinear interpolation.

**Figure 1:**
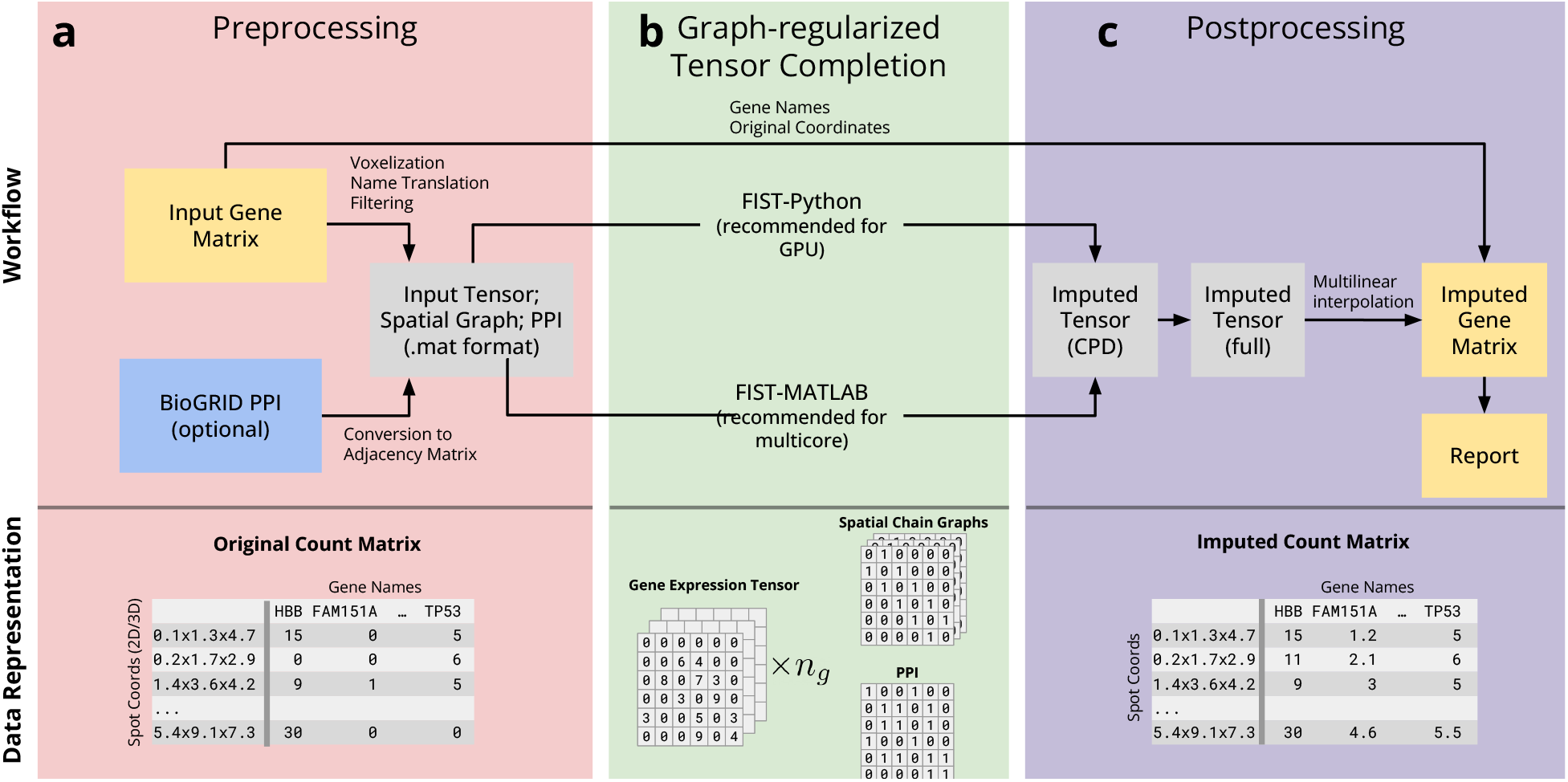
Overview of FIST-*n*D. (a) The input data is a matrix of multi-dimensional array of gene expression read counts indexed by the spatial coordinates of the spots. The preprocessing step filters and voxelizes the input data, converting it to a tensor representation by aggregating the spots. This step also constructs the spatial chain graph and a PPI graph for the next step. (b) The tensor and the graphs generated in the preprocessing step are then fed into the FIST algorithm for graph-regularized tensor completion. FIST-*n*D provides two potions, a Python implementation (for use with a GPU system) and a MATLAB implementation (for use with a multicore system). (c) The postprocessing step converts the CPD from of the tensor to the original data format through multilinear interpolation or a full tensor representation of the imputed gene expressions.

### Summary of Datasets

In the experiments, we focus on testing FIST-*n*D on four publicly available 3D spatial gene expression datasets. The first dataset is an expression atlas of the adult mouse brain (AMB) (Ortiz et al., 2020). The dataset was created by registering tissue slices sequenced using ST into a tissue atlas. The second dataset measures gene expression in the developing human heart at 6.5 PCW (DHH) (Asp et al., 2019). The third dataset measures gene expression in the human rheumatoid arthritis synovium, prepared in the same way (RAS) (Vickovic et al., 2022). The three datasets were similarly prepared through ST sequencing of parallel 2-dimensional slices, assembled to form three-dimensional data. The fourth dataset measures gene expression in a *Drosophila* embryo 16-18 hours after egg-laying (DME) (Wang et al., 2022). This dataset was created using slices of the embryo measured by high-resolution spatial sequencing technology, which was binned before being similarly arranged in the stacked slice format.

### FIST Outperforms Competing Methods

In the five-fold cross-validation, we held out a subset of the test entries in the input matrix and measure how accurately FIST with a PPI network or a diagonal gene graph reconstructs the missing entries. This cross-validation technique roughly approximates the performance of predicting unseen data. We compared FIST with other imputation algorithms, including spatial nearest-neighbor (SNN), STAGATE (Dong and Zhang, 2022), DeepImpute (DI) (Arisdakessian et al., 2019) and Markov Affinity-based Graph Imputation of Cells (MAGIC) (van Dijk et al., 2018). The SNN imputation algorithm was implemented by hand, and the provided Python packages were used to experiment with the other methods. As a major component of the graph regularization is the PPI network, we performed an ablation experiment to test whether the inclusion of the network significantly contributed to the accuracy of our results. To do this, in addition to running FIST with an organism-specific PPI downloaded from BioGrid, we also run FIST with a ‘diagonal’ gene graph, represented in adjacency matrix form as the identity matrix. For each method, we measure error by absolute error (MAE), symmetric mean absolute percentage error (SMAPE), and root mean squared error (RMSE), averaged across all genes.

Figure 2 shows the performance of FIST versus other imputation methods under the 5-fold entry-wise cross-validation. It is clear that FIST with a PPI network or a diagonal gene graph has a lower error than every other method tested, under all the three metrics. In particular, incorporation of a PPI network in the regularization also slightly improved the imputation performance by the introduction of the gene-gene relations for some of the datasets. To establish significance, we also computed 95% highest-density intervals (HDIs) for the posterior distribution of the difference between the mean MAE, SMAPE, and RMSE of FIST between FIST-*n*D with PPI against FIST-*n*D with diagonal gene graph and the other methods using the the BEST (Bayesian Estimation Supersedes the *t*-Test) method (Kruschke, 2013). Detailed results can be found in Supplementary Tables S1-S3 and Supplementary Figure S1. For two of the datasets tested (DHH, DME), FIST with a PPI significantly outperformed FIST with a diagonal graph on every metric tested.

**Figure 2:**
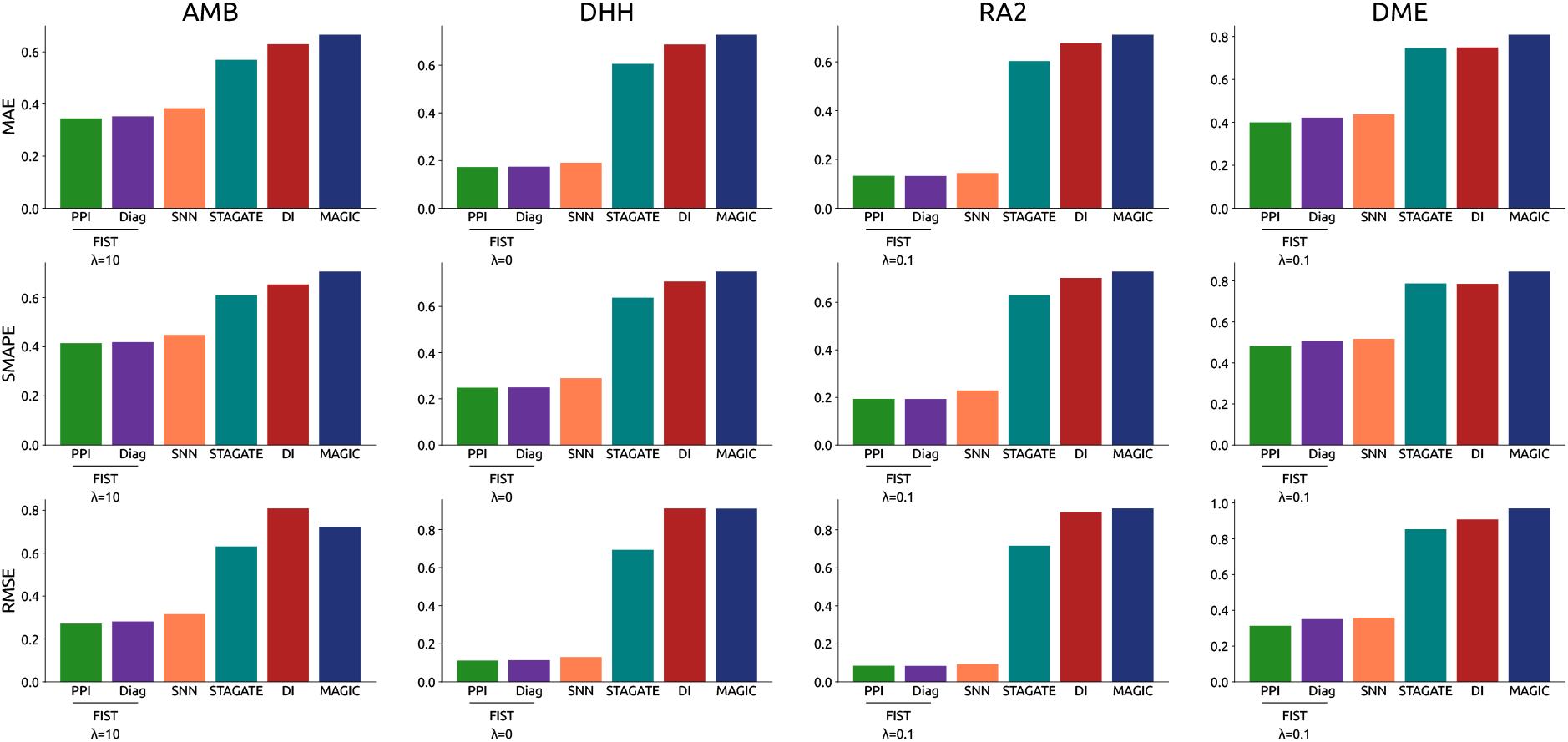
Imputation performance by five-fold cross-validation. The imputation performance was measured for each method in the four datasets. In the plots, the height of each bar corresponds to the mean value of the metric for that method, averaged over all genes.

To further characterize the error profiles of the imputation methods, Figure 3 shows a cumulative density function for the error metrics for each gene for each dataset. The errors of FIST are consistently smaller than the other imputation methods. In particular, we see that FIST does not have the concentration of high SMAPEs that we see in STAGATE, DeepImpute, and MAGIC. To examine comparisons between the datasets, we plot these data in two dimensional histograms in Supplementary Figures S2-S4, where we compare FIST with diagonal gene graph to all other methods. From these data, we conclude the FIST is the most accurate algorithm tested for imputation of stacked ST data.

**Figure 3:**
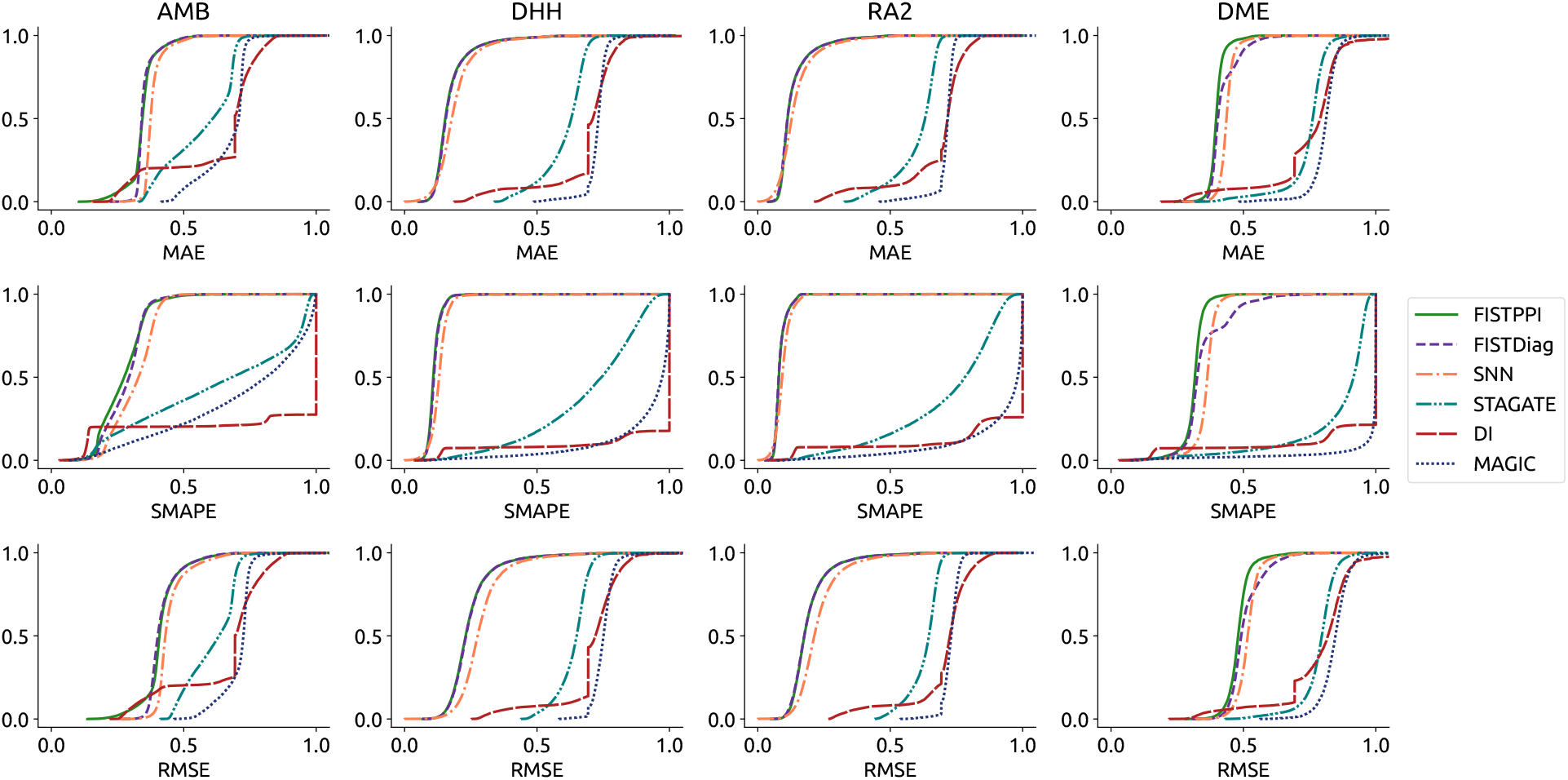
CDF of gene-wise error metrics for all the methods tested. In the plots, the x-axis is a threshold of the error measure and the y-axis is the percentage of the genes with an error lower than the threshold. Thus, the height of the distribution indicate a better performance by having a larger fraction of errors below the threshold.

### FIST Hyperparameter Tuning

To choose a hyperparameter *λ* for the regularization term of the FIST objective function, given in Equation 4, we ran a search over the set {0, 0.001, 0.01, 0.1, 1, 10}. All searches were run with a rank of 200, which is close the optimal rank reported in Li et al. (2021), and a tensor size of 20 *×* 20 *× n*_*slices*_. Plots of the error metrics of FIST models for each *λ* are shown in Figure 4. The results suggest that, in general, FIST is highly resilient to choice of the parameter *λ*.

**Figure 4:**
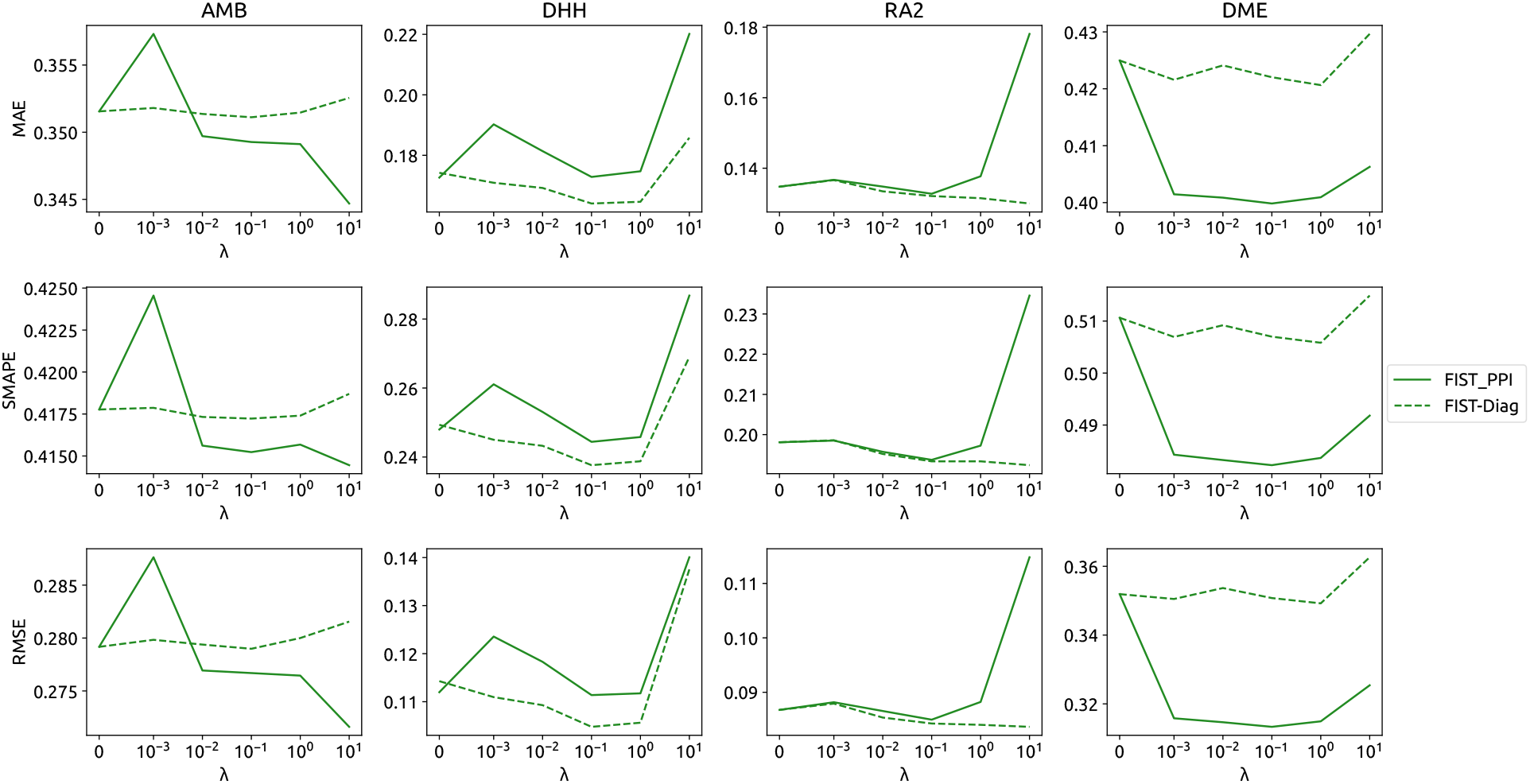
Tuning hyperparameter *λ*. The plot shows the error of FIST on each dataset under cross-validation over a set of *λ* parameters.

### Clustering Performance on GO Terms

While the previous results demonstrate that FIST accurately imputes ST data, we also validate the utility of FIST on downstream analysis tasks. Here, we consider the task of clustering genes based on their spatial profile. We performed a clustering and enrichment analysis on the data before and after imputation. For each dataset, we employ k-means clustering (*k* = 100) 10 times on gene profiles (with spatial features reduced to 1000 through PCA). We then compute the fraction of clusters enriched to some cutoff. One possible concern is the model’s ability to recover functional relevant signals without the functional information by inclusion of the PPI network, which carries functional information. To mitigate this, we tested the model’s imputation performance using a diagonal graph instead of a full PPI.

The results of the clustering analysis are shown in Figure 5. The plots show the fraction of enriched clusters as the provided *p*-value cutoff. For the human and mouse datasets, the data post-imputation has more significantly enriched clusters at almost every *p*-value between *p* = 0.001 and *p* = 0.1, suggesting that the data after imputation by FIST model is consistent with biological function. Interestingly, a lower fraction of clusters enriched is observed on the *Drosophila* dataset. However, the low performance may be indicative of lesser annotation quality in non-human organisms such as *Drosophila*. Overall, it is evident that FIST recovers gene expressions for more effective functional analysis, and with a well annotated GO database, both the accuracy of imputation and functional analysis can be improved.

**Figure 5:**
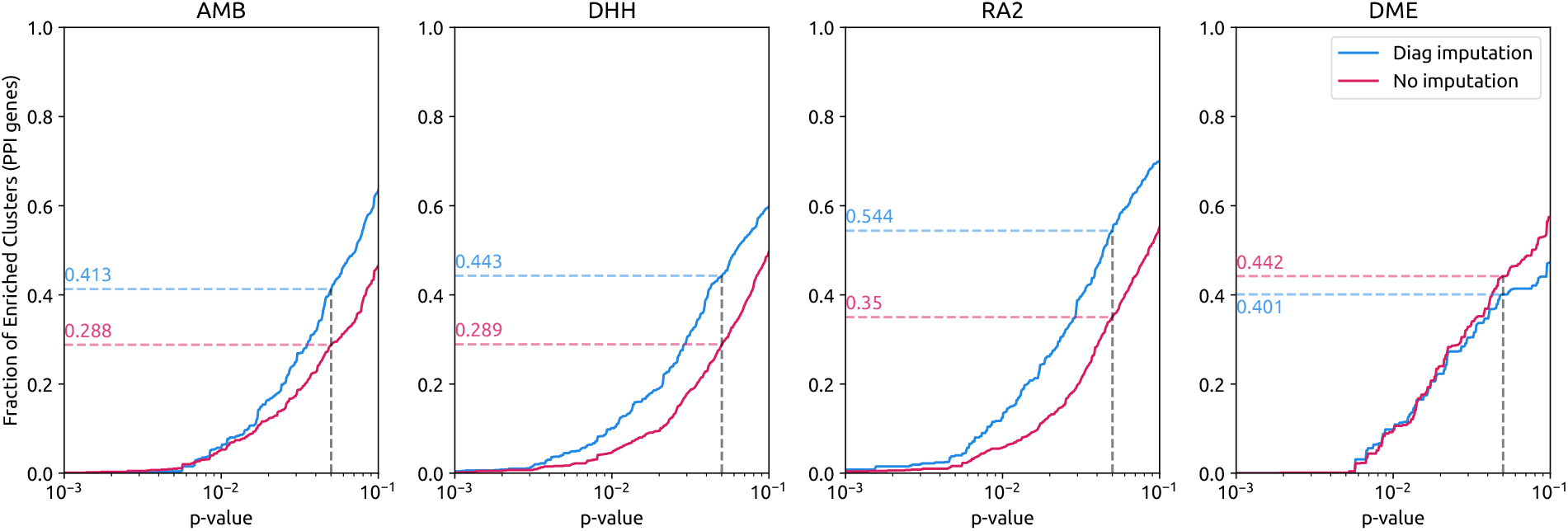
Gene ontology enrichment analysis. For each dataset, the plot shows the fraction of enriched clusters on pre- and post-imputed data. For a given *p*-value, the plots correspond to the number of clusters that had an enriched term with a smaller *p*-value. The fraction of enriched clusters at *p* = 0.05 is annotated by the dashed lines.

## Discussion

The newly developing “stacked slice” 3D spatial transcriptomics method is a promising way through which we can understand the spatial organization of tissues. Yet because these data are so sparse, we propose an imputation method, FIST, to improve downstream analyses.

Here, we provide compelling evidence that FIST-*n*D is the most accurate method for imputing 3D spatial trancriptomic data created using the “stacked slices” experimental technique. We found that FIST has significantly lower MAE, SMAPE, and RMSE when compared to other spatial and single-cell imputation methods on all four publicly available datasets tested. Furthermore, we demonstrated the utility of incorporating a PPI into the model by performing an ablation experiment.

The method not only accurately imputes 3D spatial transcriptomics data, but also improves the quality of downstream analyses. By analyzing the fraction of enriched clusters before and after imputation, we see that the imputation process leads to higher enrichment for a large range of *p*-values for the majority of the datasets tested. Furthermore, to demonstrate that this is not the result of the inclusion of PPI, we show that the result holds when the PPI network is replaced by a diagonal gene graph in the model. Thus, we have shown that FIST is both accurate and useful.

In addition to a compelling model, we also present an implementation that is fast and easy-to-use. As FIST is implemented as a command-line tool, it also is easy to use.

While we validated the method for downstream analysis using enrichment tests, an experiment comparing spatial clustering performance on labelled datasets could further validate the utility of the model. Finally, the current iteration of FIST utilizes a voxelization step to take advantage of the tensor decomposition model. Future versions that impute on the raw data instead of the voxelized data could potentially see better performance.

## Methods

### Mathematical Formulation

We begin with a review of the FIST (Fast Imputation of Spatially-Resolved transcriptomes by graph-regularized Tensor completion) method for imputation of two-dimensional ST data (Li et al., 2021), before introducing the generalization of the algorithm to FIST-*n*D. Notations are provided in Table 1.

**Table 1:** Notations.

| FIST | FIST-$nD$ | Definition |
| --- | --- | --- |
| 2 | $N$ | Number of spatial dimensions |
| $\{x, y\}$ | $\{x_1, \dots, x_N\}$ | Spatial dimensions |
| $n_i$ | $n_i$ | Size of spatial dimension $x_i$ |
| $n_p$ | $n_p$ | Size of gene dimension |
| — | $\mathcal{M} \in \mathbb{R}_+^{n_p \times n_s}$ | Original spatial gene expression spot by gene matrix |
| $\mathcal{T} \in \mathbb{R}_+^{n_p \times n_x \times n_y}$ | $\mathcal{T} \in \mathbb{R}_+^{n_p \times n_1 \times \dots \times n_N}$ | Original spatial gene expression tensor |
| $\hat{\mathcal{T}} \in \mathbb{R}_+^{n_p \times n_x \times n_y}$ | $\hat{\mathcal{T}} \in \mathbb{R}_+^{n_p \times n_1 \times \dots \times n_N}$ | Imputed spatial gene expression tensor |
| $\mathcal{Z} \in \mathbb{R}_+^{n_p \times n_x \times n_y}$ | $\mathcal{Z} \in \mathbb{R}_+^{n_p \times n_1 \times \dots \times n_N}$ | Binary mask tensor selecting nonzero entries in $\mathcal{T}$ |
| $\hat{A}_i \in \mathbb{R}^{n_i \times r}$ | $\hat{A}_i \in \mathbb{R}^{n_i \times r}$ | CPD component matrix $i$ of $\hat{\mathcal{T}}$ |
| $\mathfrak{L}(p, x, y)$ | $\mathfrak{L}(p, x_1, \dots, x_N)$ | Laplacian of PPI and spatial chain graphs |
| $\lambda$ | $\lambda$ | Regularization hyperparameter |

We first represent the spatial spots, which are arranged on a grid of size (*n*_*x*_, *n*_*y*_), and their associated spatial gene expressions as a tensor 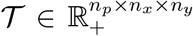 where *n*_*p*_ denotes the number of genes, so that *𝒯*_*ijk*_ is the number of reads of gene *i* at spot (*j, k*) on the slide. The learning task is to approximate *𝒯* with 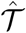 represented as a sum of outer products of vectors in the Canonical Polyadic Decomposition (CPD) form (Kolda and Bader, 2009) as follows,

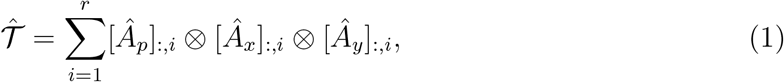

where *r* is the rank of the CPD form of 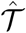 and ⊗ denotes the vector outer product. The objective function minimizes the difference between the observed and the imputed tensor under a smoothness constraint defined on the graph Laplacian of a Cartesian product of a protein-protein interaction network (PPI) with the spatial chain graphs in the *x* and *y* dimensions, denoted by 𝔏(*p, x, y*) as follows, where ⊛ represents the Hadamard product,

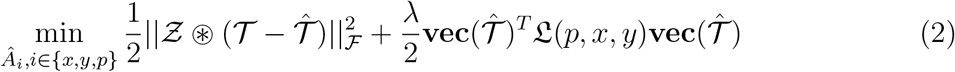

The equations from the FIST algorithm can be generalized to any number of spatial dimensions *N* following the derivations and the formulation in Li et al. (2019, 2021). Equation 1 becomes

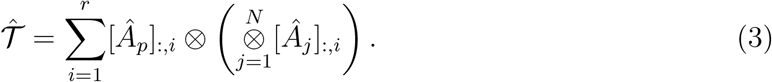

Furthermore, the objective function (Equation 2) becomes generalized as follows,

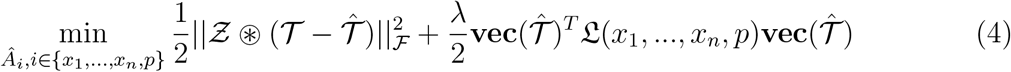

While the original authors utilize a multiplicative update rule, we use gradient descent with the Adam optimizer (Kingma and Ba, 2014). With these modifications, the FIST-*n*D algorithm is generalized to any *N* -dimensional tensor completion in theory and can be used to perform imputation on 3D spatial RNA-seq data in practice.

### FIST-*n*D Pipeline

The overall pipeline of FIST-*n*D is given in Figure 1. Each of the steps and the components are described in detail below.

FIST-*n*D is implemented as a Python package, available through the PyPI package manager, or from source at https://github.com/kuanglab/FIST-nD. Installation and running instructions and dependencies are provided on the GitHub page. The package is implemented as a command-line tool with options for inputs, outputs, PPI, hyperparameters, etc.

The pipeline starts with a multidimensional array of gene expression indexed by the spatial coordinates of the spots after mapping as input, where the first column contains the spatial coordinates and the remaining columns contain expression values of the genes. The format of the input is the same as the output provided by many existing spatial RNA-seq data processing software tools (Navarro et al., 2017) for easy integration with other pipelines.

Before converting the data to a tensor format, a few preprocessing steps need to be applied. FIST-*n*D first attempts to determine if one of the axes is composed of discrete slices so that these slices are represented as consecutive tensor indices in that dimension. The tool optionally allows the user to rotate the data using a PCA transformation in order to have the dimensions with the highest variance line up with the tensor axes for better performance after voxelization.

If the user is working with 2-dimensional 10X Visium data, the data are already arranged in a spatially uniform grid for tensor completion. However, if we have data with spots in continuous space, we must represent these data as a *N* -way tensor for operating on a tensor in the CPD form. The process of converting points in continuous space to a tensor (akin to voxelization) for the FIST input is as follows, shown graphically in two dimensions in Supplementary Figure S5. Voxel boundaries are determined by taking evenly spaced quantiles of the distribution of spots along each spatial dimension. In the binning process, let *s* index the spots as (*x*_*s*_, *y*_*s*_, *z*_*s*_) with expression vectors *g*_*i*_, we calculate *𝒯*_:,*ijk*_, each (*i, j, k*)-th gene fiber in the tensor by the following metric:

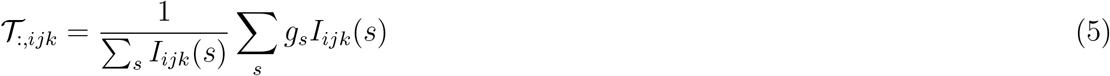

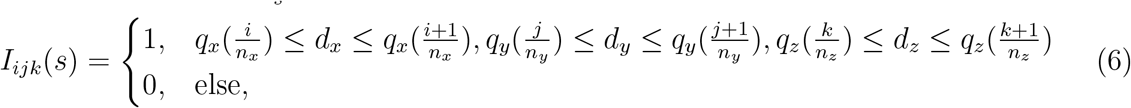

where *q*_*d*_(.) is the quantile function for dimension *d* ∈ *{x, y, z}*. Here, *I*_*ijk*_(*s*) is an indicator if spot *s* is a member of voxel (*i, j, k*). After the voxelization, any voxel with less than three total read counts is set to zero and all the genes with 8 or less total counts after this step are also removed from the tensor. Finally, a log-transform is then applied to the voxelized tensor.

We optionally allow the user to specify a PPI network as input for the graph regularization, which can be downloaded from BioGRID (https://downloads.thebiogrid.org/BioGRID/Release-Archive/BIOGRID-4.4.201/) (Oughtred et al., 2021). BioGRID contains PPI networks for a variety of organisms, represented as a list of pairs of interacting genes. Our software converts this representation to an adjacency-matrix format, using the MyGene annotation service to match the gene names provided by the user in the original count matrix to their official gene symbols (Wu et al., 2013).

If no PPI is provided by the user, FIST-*n*D by default uses a dummy diagonal graph, where every gene interacts only with itself. This has the effect that the model only uses the spatial information for smoothing. This default construction of FIST-*n*D outperforms other competing imputation methods in the crossvalidation experiments (see Results section), although the imputation result may be better when a real PPI is used.

The optimization algorithm stops when a user-specified maximum number of iterations for the algorithm to run is reached. After this, the algorithm will choose the imputation from the iteration that contained the lowest test error on a held-out validation set.

After the completion of the optimization, the CPD form of the imputed tensor can be utilized for conversion back to the original format, a matrix of gene expressions by spots. A multilinear interpolation algorithm is used for the conversion. If (*x, y, z*) is in voxel (*i, j, k*) with corner (*v*_*i*_, *v*_*j*_, *v*_*k*_), 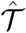 is the imputed tensor, and 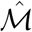 is the imputed spot by gene matrix, then

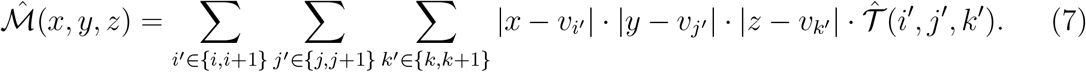

This function considers all neighboring voxels of the spot to allows further variation in read counts to occur within voxels, reducing the importance of binning size as a hyperparameter.

The final output of the imputed data is in the same format as the original input data, along with an optional report of the imputation. This report includes parameter details, plots of validation error, a list of the genes with the highest variance across all spots in the post-imputation data, and side-by-side plots of selected genes pre-imputation and post-imputation. Furthermore, we provide a separate script for the side-by-side visualization of genes pre- and post-imputation in an interactive 3D plot.

### Evaluation and Crossvalidation

#### Five-fold Crossvalidation and Error Metrics

To test the methods, we performed 5-fold entry-wise crossvalidation by holding out a subset of the entries (20%) from the original gene expression matrix. This modified input matrix with 80% training entries was run through our pipeline, and the imputed entries were compared to the original held-out test entries (both log-transformed). For each gene *g*, we evaluated the following three metrics:

- Mean absolute error (MAE): 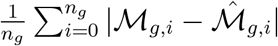.
- Symmetric mean absolute percentage error (SMAPE): 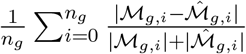.
- Root mean squared error (RMSE): 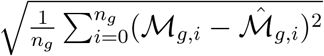.

Then, the overall performance is evaluated by the mean and variance of these metrics over the genes. Note that for all the metrics, nonzero entries in the original expression matrix *ℳ* were considered as unknowns and not used in the calculating the errors.

#### Related Methods for Comparison

FIST-*n*D was compared with four existing imputation methods, described below,

- **SpatialNN:** For a baseline method that also incorporates spatial information, we implemented a spatial single nearest-neighbor algorithm that takes the voxelized tensor as input and imputes missing tensor entries based on the closest tensor entry (as measured by Euclidian distance between tensor indices).
- **STAGATE (Spatially resolved Transcriptomics with an Adaptive Graph ATtention auto-Encoder):** STAGATE implements a graph attention-based autoencoder for spatial transcriptomic embedding and clustering (Dong and Zhang, 2022). From the associated decoder, we recover the model’s imputations of 3D stacked slice ST data.
- **DeepImpute:** DeepImpute imputes scRNA-seq data by dividing genes into correlated classes, each of which are imputed by a sub-neural network (Arisdakessian et al., 2019). However, this model does not take into account spatial information, being originally developed for single-cell data.
- **MAGIC (Markov Affinity-based Graph Imputation of Cells):** Another scRNAseq imputation technique, MAGIC imputes single cell RNA-seq data by constructing a Markov distance matrix for random walk (van Dijk et al., 2018). This method has consistently ranked among the top single cell RNA-seq data imputation methods (Hou et al., 2020).

**Table 2:** Summary of 3D spatial transcriptomics datasets. Density is calculated as number of nonzero entries in the gene expression matrix divided by the total number of entries in the gene expression matrix.

| Dataset | # of Genes | # of Spots | Slices | Density |
| --- | --- | --- | --- | --- |
| AMB | 23371 | 34103 | 40 | 0.1892 |
| DHH | 19336 | 1467 | 9 | 0.1092 |
| RAS | 12333 | 2338 | 7 | 0.1456 |
| DME | 13668 | 15295 | 16 | 0.0464 |

### Expression Clustering Enrichment Analysis

To validate the ability of FIST-*n*D to extract biological truth, we imputed all the datasets and tested the extent to which the imputation improved gene clustering performance, as measured by GO enrichment. The post imputation matrix was defined as 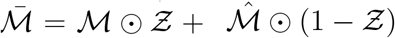, where 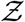 is the binary mask matrix, the pre-imputation matrix was given by *ℳ* and the output of FIST is given by 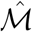. Clustering was done by first reducing the number of features (spots) for each gene down to 1000 with PCA and then applying k-means with *k* = 100 (Pedregosa et al., 2011). This was repeated 10 times with different random seed initializations to reduce variability by local minima (Macqueen, 1967). Enrichment analysis on each cluster was performed with the enrichGO function in the R clusterProfiler package with FDR correction applied (Wu et al., 2021). The size of 20 by 20 by 20 and a value of *λ* = 1 was used for the imputation by FIST-*n*D models. No PPI was used to prevent leakage of functional information.

### 3D Spatial Transcriptomics Datasets

Four 3D spatial gene expression datasets including three ST datasets and one high-resolution Stereo-seq dataset are analyzed in the experiments.

- Adult Mouse Brain (AMB): This dataset was collected from a sample of adult mouse brain (AMB) tissue (Ortiz et al., 2020). The authors took 40 parallel slices along the anteroposterior axis of the tissue, which were then registered to produce three dimensional coordinates for each spot. The data were downloaded from the authors’ website (https://www.molecularatlas.org/download-data). The meta table and the raw expression matrix were downloaded from the processed data section, which were combined to create the input file.
- Developing Human Heart (DHH): This dataset was collected from a sample of developing human heart (DHH) tissue taken 6.5 weeks post-conception (Asp et al., 2019). The authors obtained 9 parallel slices of the tissue sample with 105-219 spots per slice covered by the tissue (of 1,007 maximum possible). The data were downloaded from the authors’ GitHub repo (https://github.com/MickanAsp/Developmental_heart). The data were then processed to combine the atlas and gene tables into one table in the format described, using the spot.pos column as an index.
- Rheumatoid Arthritis Synovium (RA2): This dataset was collected from a rheumatoid arthritis synovium tissue sample (Vickovic et al., 2022). From the six patient samples presented in the original study, we took the dataset from the second patient (RA2), as it contains 7 slices, with the most number of slices among all six samples. The data were downloaded from the repository Single Cell Portal (https://singlecell.broadinstitute.org/single_cell/study/SCP1414/3dst-ra). Because the separate slices were provided as separate files, the location and position information for each slice was combined, with all the slices being concatenated into a single dataset.
- Drosophila Melanogaster Embryo (DME): This dataset was collected from a Drosophila embryo at 16-18 hours after egg-laying and contains 16 slices (Wang et al., 2022). The data were collected via Stereo-seq, and binned into 20×20 bins of DNBs. The data are available at https://db.cngb.org/stomics/flysta3d/download.html.The dataset was rotated using a PCA transfomation of the x and y axes so that the major axis of the ellipsoid was parallel to the x axis, so that the shape fit more efficiently for binning.

## Availability and Requirements

The data used for the project are publicly available datasets published elsewhere. They can be found at the links in the **3D Spatial Transcriptomics Datasets** section.

The tool is available from PyPI at https://pypi.org/project/fistnd/ or from source at https://github.com/kuanglab/FIST-nD.

## Acknowledgements

The authors acknowledge the Minnesota Supercomputing Institute (MSI) at the University of Minnesota for providing resources that contributed to the research results reported within this paper (URL: http://www.msi.umn.edu).

## Supplementary Information

**Table S1:** 95% highest density intervals for posterior distribution of the difference between mean MAE for genes imputed by FIST versus competing methods.

| Dataset | FIST-Diag | SNN | STAGATE | DeepImpute | MAGIC |
| --- | --- | --- | --- | --- | --- |
| AMB | (-0.004,-0.003) | (0.030,0.031) | (0.246,0.251) | (0.354,0.358) | (0.351,0.353) |
| DHH | (0.001,0.003) | (0.023,0.025) | (0.470,0.473) | (0.582,0.584) | (0.577,0.579) |
| RA2 | (-0.001,0.000) | (0.014,0.016) | (0.514,0.517) | (0.606,0.608) | (0.604,0.605) |
| DME | (0.007,0.009) | (0.038,0.040) | (0.365,0.366) | (0.398,0.401) | (0.415,0.417) |

**Table S2:** 95% highest density intervals for posterior distribution of the difference between mean SMAPE for genes imputed by FIST versus competing methods.

| Dataset | FIST-Diag | SNN | STAGATE | DeepImpute | MAGIC |
| --- | --- | --- | --- | --- | --- |
| AMB | (0.009,0.012) | (0.043,0.046) | (0.352,0.361) | (0.481,0.492) | (0.459,0.469) |
| DHH | (0.002,0.003) | (0.021,0.022) | (0.606,0.613) | (0.891,0.892) | (0.857,0.860) |
| RA2 | (-0.001,-0.000) | (0.011,0.013) | (0.685,0.692) | (0.920,0.921) | (0.895,0.897) |
| DME | (0.009,0.011) | (0.046,0.048) | (0.594,0.597) | (0.683,0.683) | (0.681,0.682) |

**Table S3:** 95% highest density intervals for posterior distribution of the difference between mean RMSE for genes imputed by FIST versus competing methods.

| Dataset | FIST-Diag | SNN | STAGATE | DeepImpute | MAGIC |
| --- | --- | --- | --- | --- | --- |
| AMB | (-0.009,-0.007) | (0.020,0.022) | (0.205,0.209) | (0.302,0.307) | (0.307,0.308) |
| DHH | (0.001,0.004) | (0.044,0.047) | (0.415,0.417) | (0.519,0.522) | (0.519,0.521) |
| RA2 | (-0.002,0.001) | (0.036,0.039) | (0.467,0.470) | (0.553,0.555) | (0.552,0.554) |
| DME | (0.013,0.015) | (0.035,0.037) | (0.315,0.318) | (0.349,0.351) | (0.366,0.369) |

**Figure S1:**
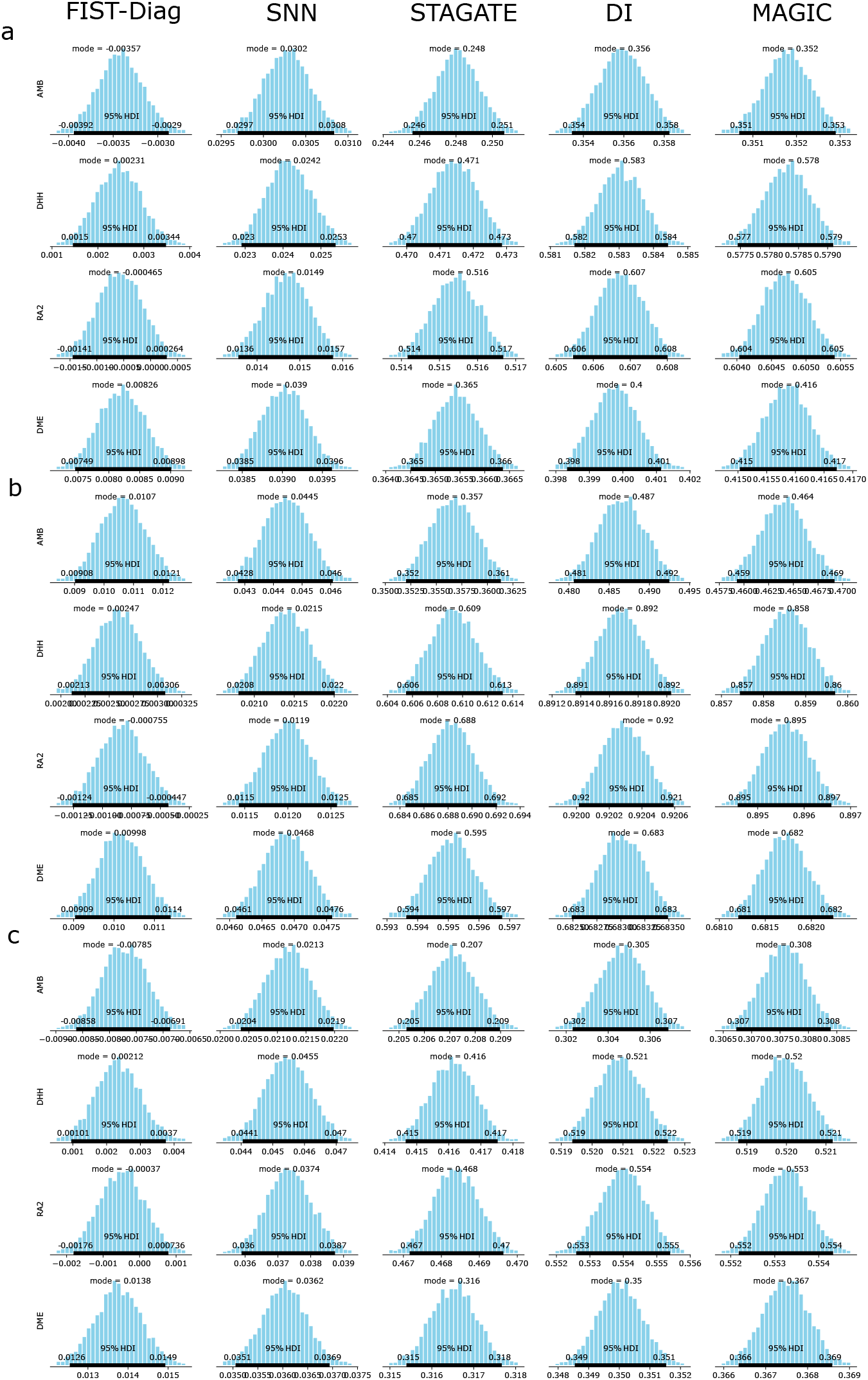
Posterior distributions of differences between mean (a) MAE, (b) SMAPE, and (c) RMSE between FIST and other methods. From left to right, FIST (diagonal PPI), SNN, STAGATE, DeepImpute, and MAGIC. The 95% HDI is highlighted. Computed using BEST (Bayesian Estimation Supersedes the *t*-Test).

**Figure S2:**
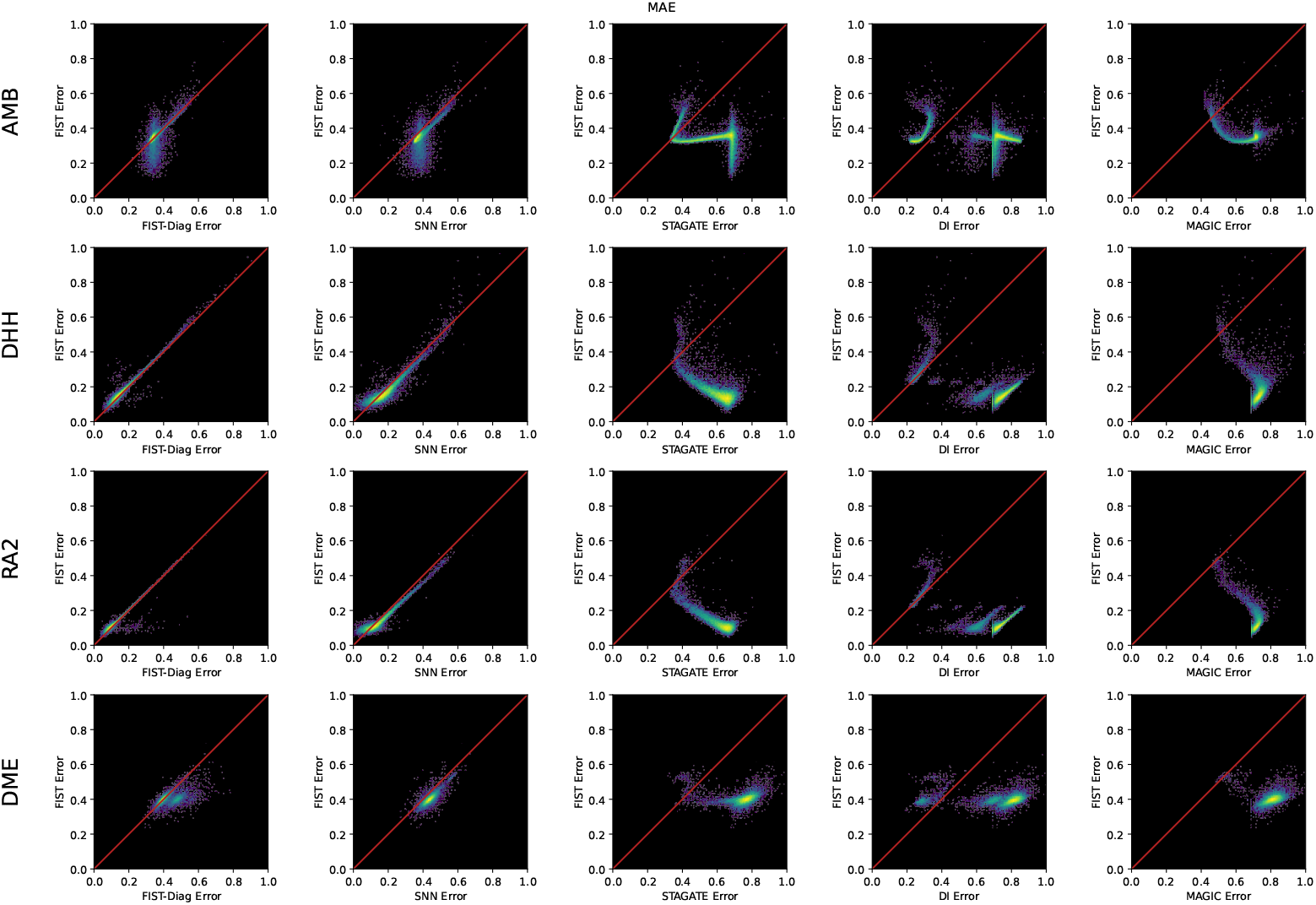
2D histogram of MAEs for each gene for FIST against each method.

**Figure S3:**
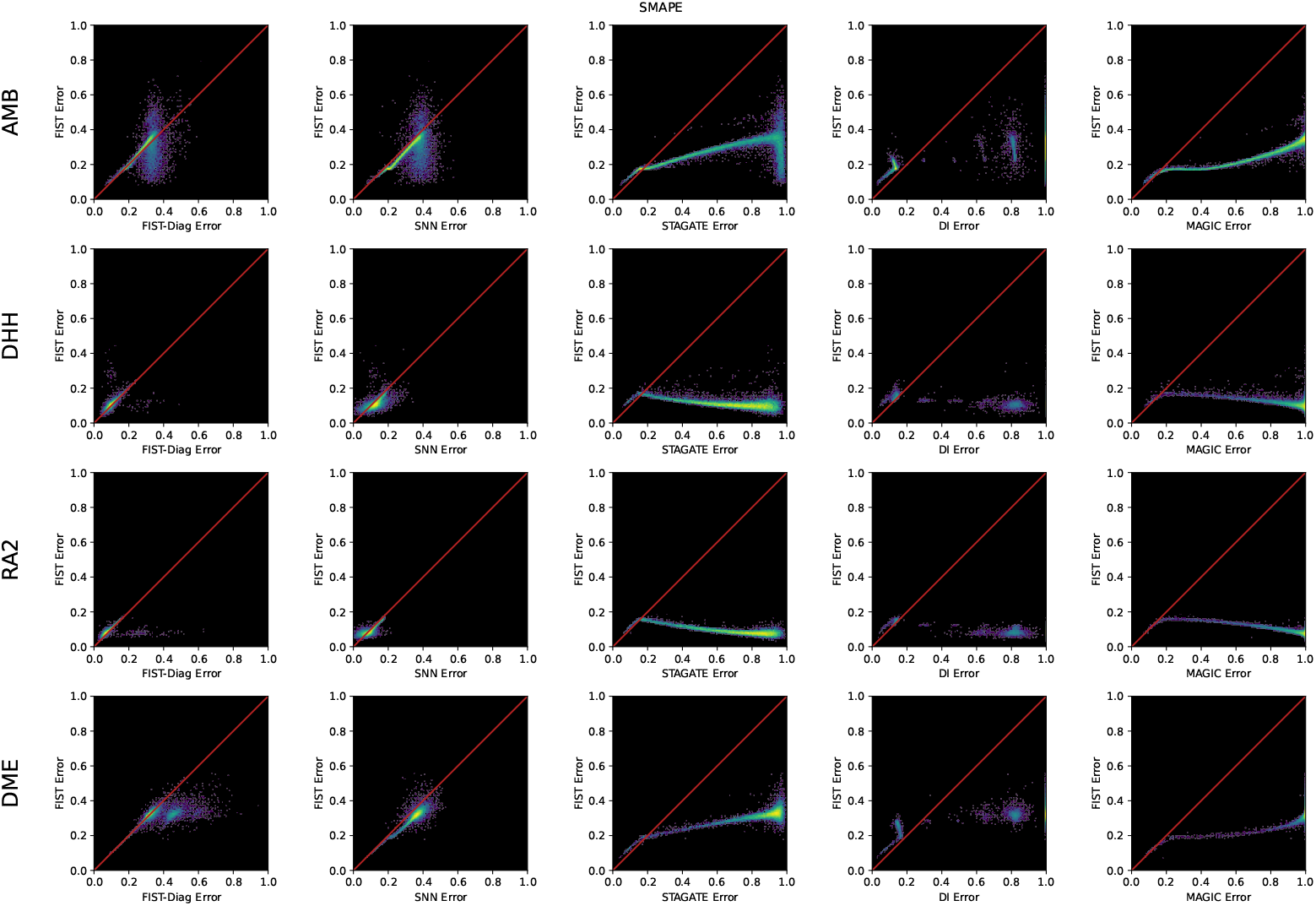
2D histogram of SMAPEs for each gene for FIST against each method.

**Figure S4:**
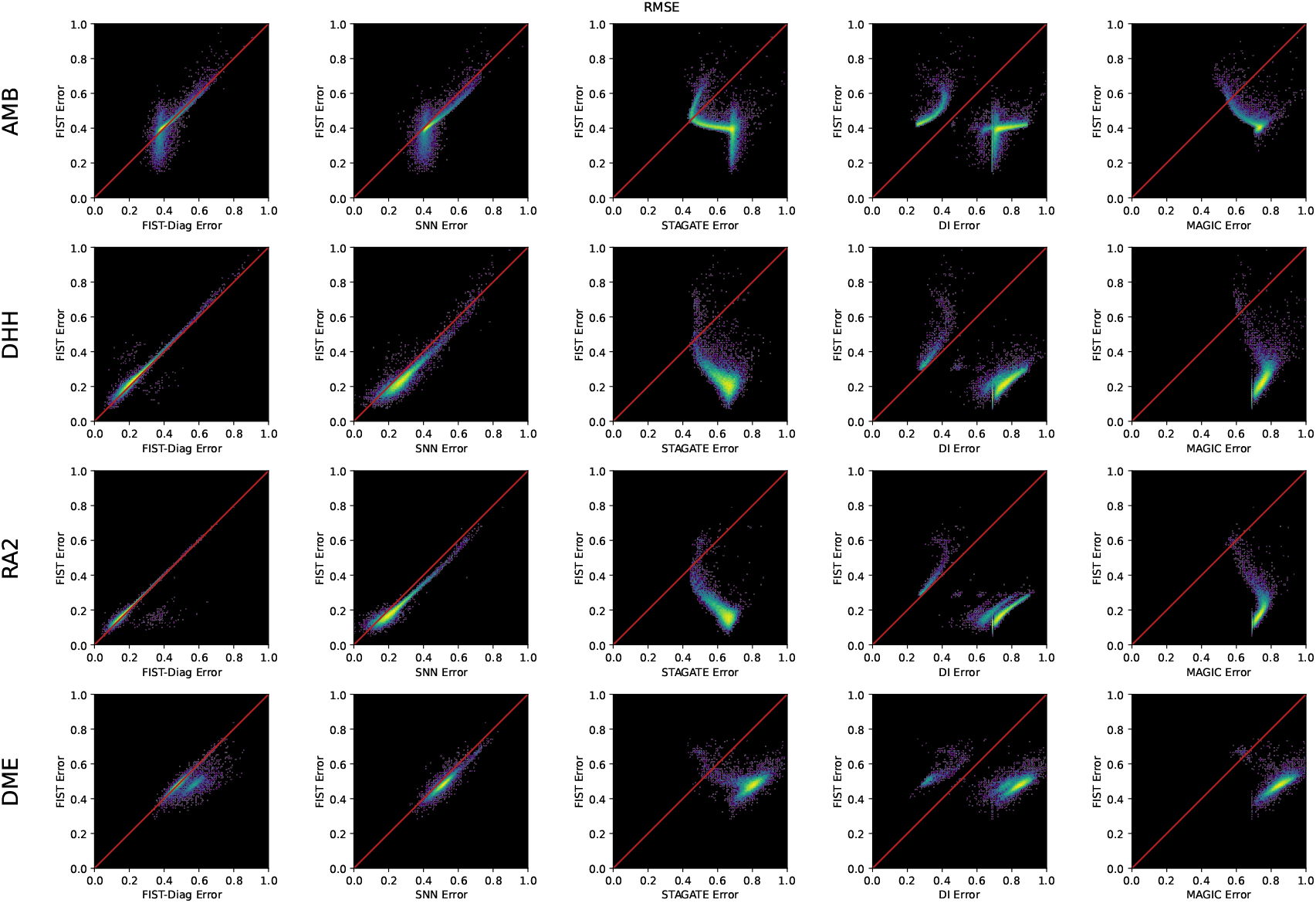
2D histogram of RMSEs for each gene for FIST against each method.

**Figure S5:**
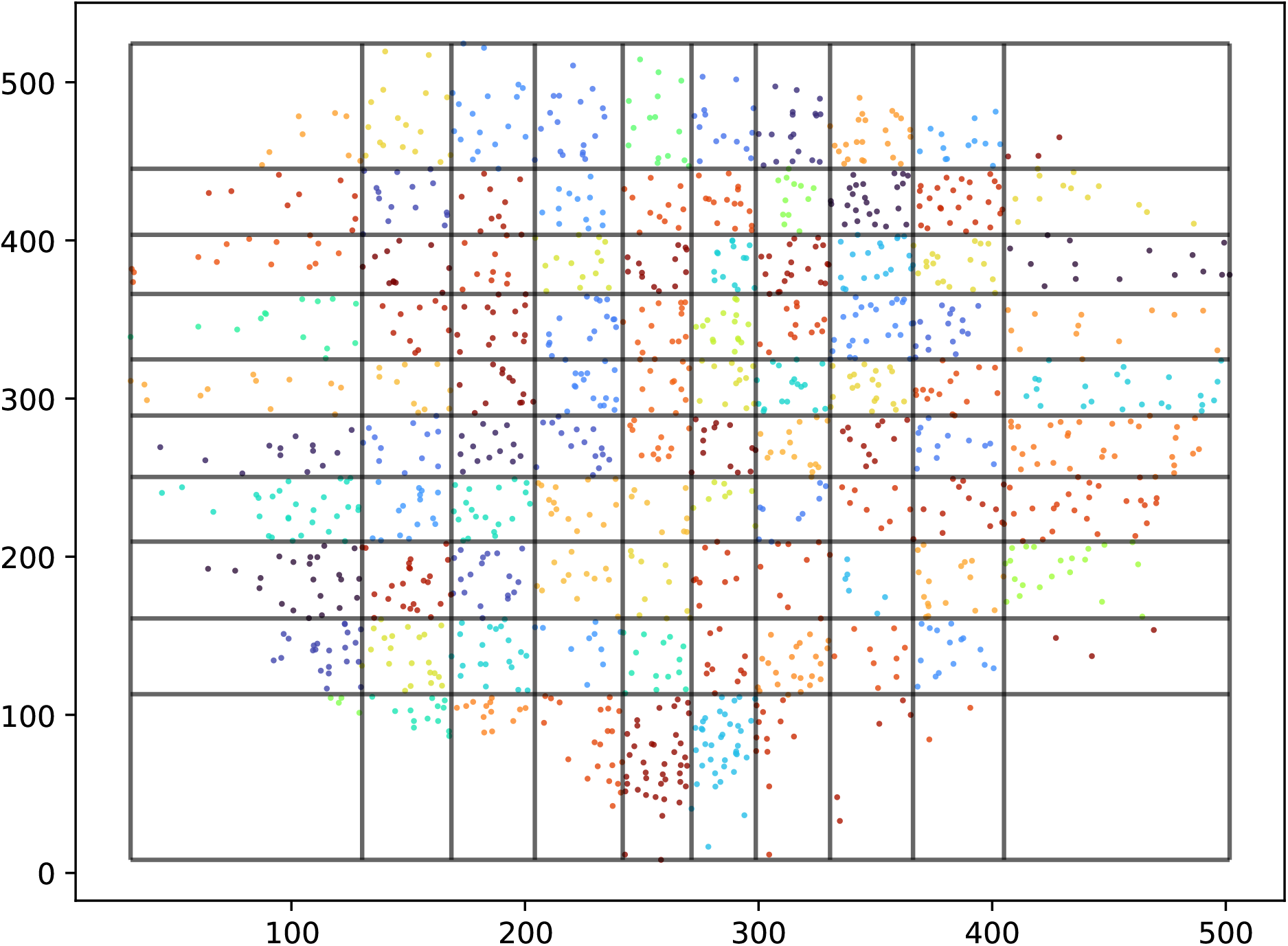
Visual representation of the binning process in two dimensions. Voxel boundaries are drawn on each axis corresponding to equally spaced quantiles of the data. The tensor entries for each gene are determined by the mean of all gene counts of spots that fall in the bin (here represented by color).

